# Predicting Cortical Bone Resorption of Mouse Tibia in Disuse Condition Caused by Transient Muscle Paralysis

**DOI:** 10.1101/2023.11.21.568078

**Authors:** Himanshu Shekhar, Sanjay Singh, Jitendra Prasad

## Abstract

Load removal from the load-bearing bone, such as in the case of extended space travel and prolonged bed rest, harms bone health and leads to severe bone loss. However, the constitutive idea relating the quantity of bone loss to the absence of physiological loading is poorly understood. This work attempts to develop a mathematical model that predicts cortical bone loss at three sections: ‘distal,’ ‘mid-section,’ and ‘proximal’ along the length of a mouse tibia. Load-induced interstitial fluid flow-based dissipation energy density has been adopted as a stimulus to trigger mechanotransduction. The developed model takes the loss of stimulus due to the disuse of bone as an input and predicts the quantity of bone loss with spatial accuracy. We hypothesized that the bone loss site would be the site of maximum stimulus loss due to disuse.

To test the hypothesis, we calculated stimulus loss, i.e., loss of dissipation energy density due to bone disuse, based on the poroelastic analysis of the bone using a finite element method. A novel mathematical model has been then developed that successfully relates this loss of stimulus to the in-vivo bone loss data in the literature. According to the developed model, the site-specific mineral rate is found to be proportional to the square root of the loss of dissipation energy density. To the author’s best knowledge, this model is the first of its kind to compute site-specific bone loss. The developed model can be extended to predict bone loss due to other disuse conditions such as long space travel, prolonged bed rest, etc.

## 1. Introduction

The bone is an example of an optimized structure. The internal architecture of a bone is adapted to the external mechanical load exerted during daily physical activity ^1 2 3 4^. However, certain conditions, such as extended space travel ^5 6^, prolonged bed rest ^7 8^, or lower-limb paralysis due to spinal cord injury^9^, disrupt these physical activities. A reduction in mechanical load on the weight-bearing bone under these conditions leads to significant bone loss ^10^. Recent studies have shown that the microgravity environment in spaceflight is so harmful that even one year after the mission, the lost bone tissue is not entirely recovered, possibly leading to osteoporosis ^11^. For example, data from the Russian Mir and the International Space Station reveals that crew members of long-duration space flights lose approximately 1.2-1.5% bone mass per month in their proximal femur ^12^, and a six-month mission results in a 2% loss in cortical density in the tibial diaphysis. However, spaceflight studies are prohibitively expensive. Therefore, partial weight bearing ^1314^ and hindlimb unloading(via tail-suspension) were used to simulate such a study ^15^. For example, 14 days of hind-limb unloading causes significant bone loss on the endocortical surface of the femoral diaphysis ^16^. Botox has also been explored as a method for muscle dysfunction^17^, which leads to the disuse of bone and, thus, bone loss ^1819^. This method has shown that Botox-induced endo-cortical bone loss is heterogeneous along the tibial length ^20^. Despite these observations, the cellular and molecular mechanism linking the loss of mechanical loading to bone tissue deterioration remains poorly understood.

It is well established that osteocytes function as mechanosensory cell ^21 22^ and they are intricately within the bone matrix via the lacuno-canalicular network (LCN). Experimental studies suggest that load-induced fluid flow within the lacuna-canalicular system (LCS) plays a critical role in transporting nutrients and enhancing osteocyte metabolic function, which is essential for maintaining bone health ^23^. Additionally, load-induced fluid flow is recognized as a key stimulus for bone mechanotransduction^24 23 25 26^. Research also indicates that this fluid flow through LCS creates shear stresses on the cell wall ^27^ which activates the mechano-sensing cell osteocytes. Tracer techniques have further confirmed the presence of fluid flow within the LCS ^28^, although measuring fluid velocity experimentally is challenging due to the inaccessibility of the LCN. Only a few studies have successfully measured fluid velocity ^29^, which is why the finite element method is commonly employed to estimate it ^30 31 32,33 34 35 36^. Recent studies have demonstrated that Botox-induced muscle paralysis significantly alters bone morphology ^37^, resulting in decreased fluid velocity around the osteocytes ^38^. This finding has inspired the current work, which models bone loss by considering the reduction of load-induced fluid flow.

When bone is mechanically strained during physiological loading, a pressure gradient develops within the lacuna-canalicular network (LCN), causing fluid to flow through the canalicular channels (Fig 1). Due to the fluid’s viscosity, this flow generates shear stress on the cell membrane and leads to energy dissipation. The dissipated energy, which is a function of fluid velocity, is assumed to be a stimulus to the mechano-sensing cell, i.e., osteocyte ^34^. Studies indicate that in loaded bone, the resultant fluid velocity is higher at the endocortical surface but negligible at the periosteal surface ^39^ due to the permeable and impermeable nature of the respective surfaces ^40^. When Botox-induced calf muscle paralysis pauses the locomotion, bone strains drop to zero, and fluid flow through the porosities of LCN ceases (Fig 1(d)). Consequently, there is a more significant loss of fluid velocity at the endocortical surface, which correlates with the in-vivo pattern of bone loss observed at the endocortical surface following Botox-induced muscle paralysis.

**Fig. 1.**
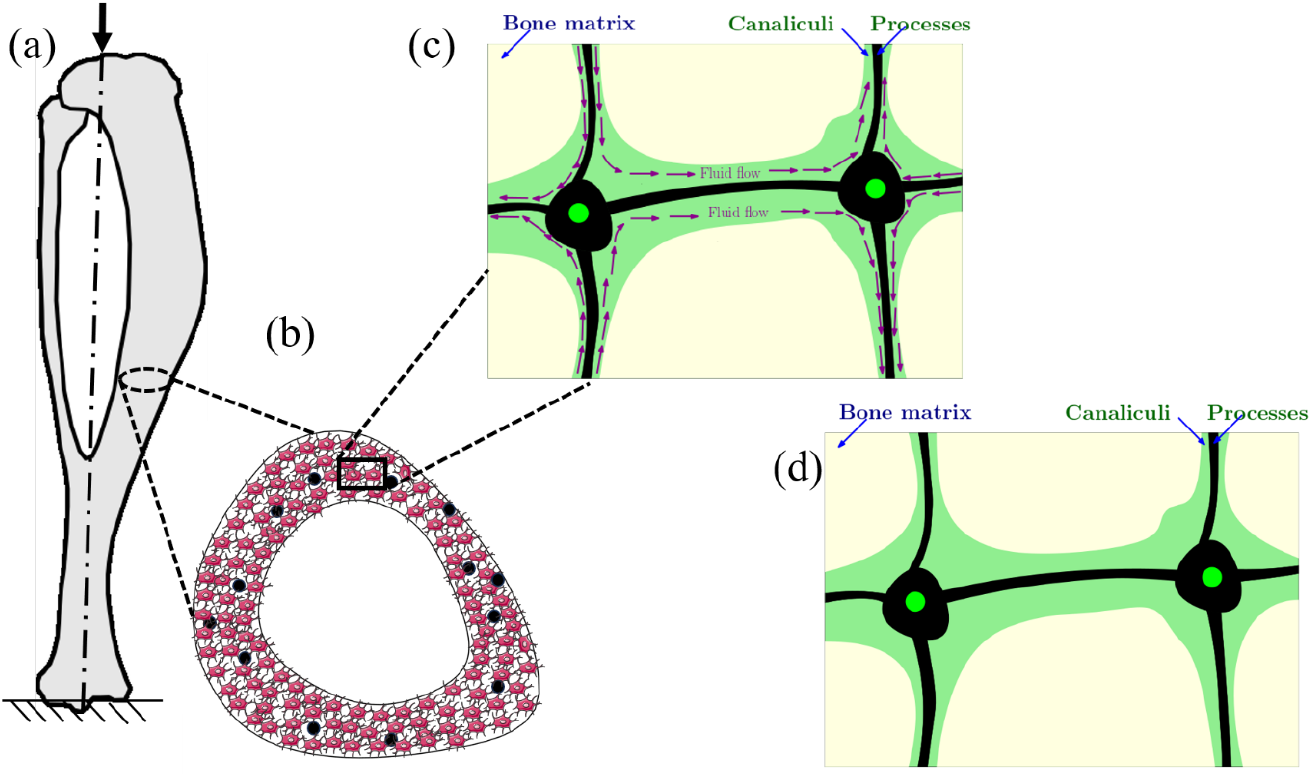
Comparison of fluid flow within the LCN porosity between a physiologically loaded and disused tibia: (a) axially loaded tibia, (b) schematic of the osteocyte network at a cross-section of interest (black filled circles represents cortical canal), (c) fluid flow within the LCS of a physiologically loaded tibia (indicated by arrows showing fluid movement), and (d) absence of fluid flow within the LCS in the case of bone disuse.

The study focuses on the loss of mechanical stimulus, specifically the reduction in fluid flow-based dissipation energy density due to bone disuse, and introduces a mathematical model to predict cortical bone loss accurately. The developed model was validated using published data on Botox-induced bone loss ^20^. Statistical tests were conducted to assess the model’s accuracy in predicting in-vivo results. The model can be extended on the availability of relevant experiment data to predict remaining disuse conditions such as long space travels, prolonged bed rest, etc. The paper is structured as follows: Section 2 outlines the adopted methodology, Section 3 presents the results, Section 4 discusses the findings, and Section 5 provides the conclusions.

## 2. Methods

### 2.1 Theory of Poroelasticity

Mouse tibia was considered a linear isotropic porous material consisting of a solid matrix and fluid-saturated pores. The constitutive equations for such material are as follows ^41^.

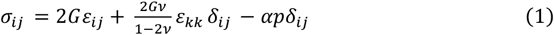

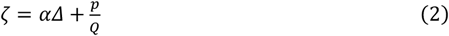

where *σ*_*ij*_ and *ϵ_ij_* are stress and strain tensors, respectively (*i* or *j* represents the Cartesian coordinates, i.e., *i* or j=1,2,3 corresponds to the x, y, z coordinates). *G* and *ν* are the modulus of rigidity and Poisson’s ratio, respectively. *p* is the interstitial pore pressure, *ζ* is the increment of the water volume per unit volume of the solid matrix, *δ_ij_* is the Kronecker delta (i.e., *δ_ij_* = 1 if *i*=*j*; *δ_ij_* = 0 otherwise), *Δ* is the dilation that represents the volumetric strain developed in the pore, α, and 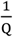 are the constants.

Equation (3) describes the fluid flow rate through a porous media following Darcy’s law:

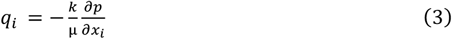

where, *q*_i_ is the fluid mass flow rate, while *k* and *µ* represent the isotropic intrinsic permeability and dynamic viscosity of the fluid, respectively.

The mass balance equation is as follows:

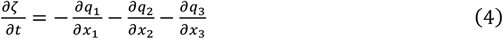

The above equations finally lead to the following differential equation that needs to be solved:

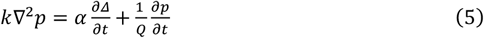

Additionally, the linear momentum balance can be written as

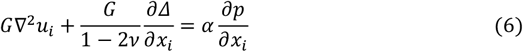

Where *u*_*i*_ is the displacement of the solid matrix. The above Eqs. (5) and (6) together have 4 unknowns to be solved for – 3 displacements components (viz. *u*_1_, *u*_2_ and *u*_3_) and the fluid pore pressure *p*. The problem was solved using a finite element commercial ABAQUS software^42^, as detailed in section 2.6.

### 2.2 Cage Activity of a Mouse

The cage activity data for the mice were taken directly from the earlier study by Hasriadi et al., where six mice were monitored over a 24-hour period, and their activities were recorded ^43^.

The typical activities observed in the cage included locomotion, rearing, immobilization, and climbing:

- **Locomotion**: Locomotion was the most common activity. During the locomotion, the tibia experiences ground reaction forces, which can be resolved into axial compression and transverse shear forces ^44^. It was found that the magnitude of shear force is very minimal compared to the axial component and could therefore be ignored ^45 38^. As a result, the primary force acting on the mouse tibia during locomotion is axial compression.
- **Rearing**: Rearing was the second most frequent activity. When rearing, the mouse stands on its hind limbs, subjecting the tibia to pure axial loading.
- **Immobilization**: During immobilization, static loading is applied to the tibia. However, it is well established that static loading does not promote new bone formation^46^.
- **Climbing**: Climbing leads to no ground reaction forces on the mouse tibia.

These observations suggest that the axial loading is typically significant during the cage activities of a mouse.

### 2.3 Bone geometry

The tibia of a 16-weeks-old C57BL6J mouse was scanned with a 21µm-voxel size, and micro-CT images were obtained ^44^. The raw micro-CT images were processed using appropriate tools, including flip, rotate, resize, crop, and black and white. An appropriate threshold was chosen to capture the tibial cross-section. All processing steps were carried out using the in-house software “Priffer”, as illustrated in Fig. 2. The processed micro-CT images were then rendered to generate a voxel-based 3D model of the tibia for finite element analysis.

**Fig. 2.**
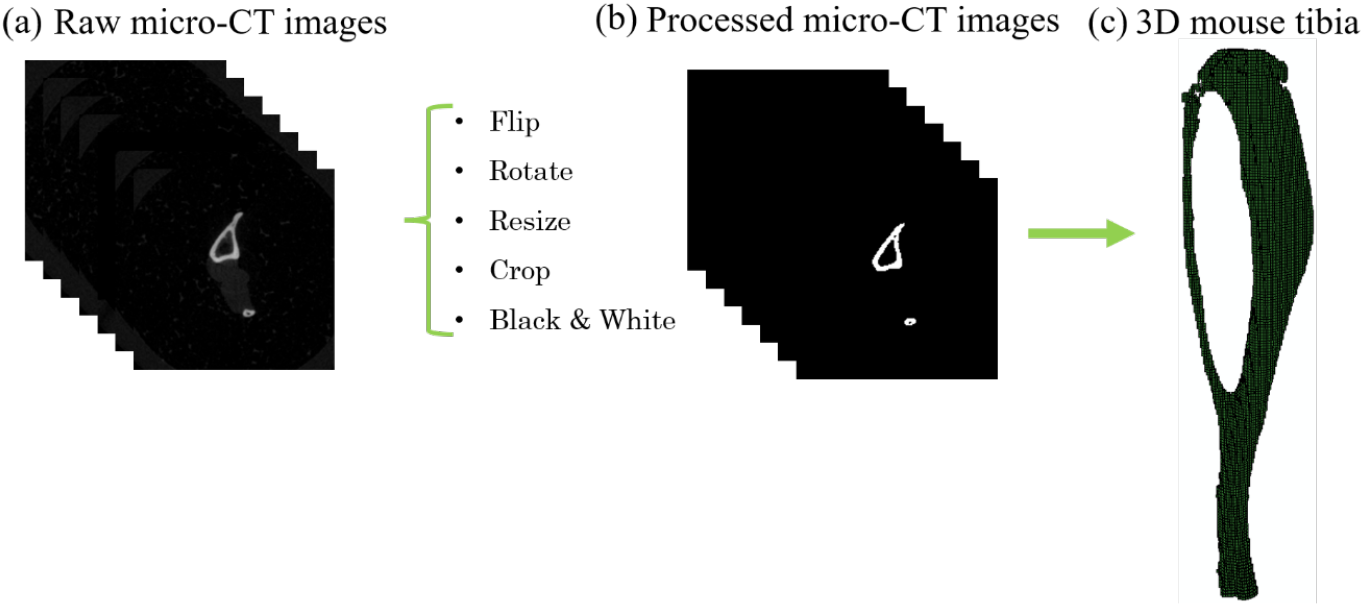
Construction of a voxel-based tibia model: (a) a set of raw micro-CT images of a 16-week mouse tibia, (b) processed micro-CT, raw images were processed using the needed tools such as Flip, rotate, resize, crop, and Black and white. An appropriate threshold was chosen to capture the bone cross-section (c) rendered 3D mouse tibia having a brick element(hexahedral element) for finite element analysis.

### 2.4 Porosity and permeability of cortical bone

The cortical bone has porosities of different length scales, such as vascular and lacune-canalicular porosity, which influence bone permeability. It is widely accepted in the literature that the loading-induced fluid flow inside the LCN porosities plays an important role in bone mechanotransduction. Accordingly, only LCN porosity has been considered, viz. 3.79 × 10^−21^ *m*^247^, as also presented in Table 1.

**Table 1:** Poroelastic material properties used in the simulations adapted from ^34^ except for the permeability value ^47^.

| Properties | Magnitude | Unit |
| --- | --- | --- |
| Young's modulus of bone | 12 | GPa |
| Fluid bulk modulus | 2.3 | GPa |
| Solid bulk modulus | 17 | GPa |
| Porosity | 0.05 |  |
| Poisson's Ration | 0.3 |  |
| Intrinsic permeability (k) | $3.79 \times 10^{-21}$ | $m^2$ |

### 2.5 Loading and boundary conditions

The physiological loading of a caged mouse is dominated by axial loading (Fig 3(a)). To mimic the physiological condition, a Haversine loading waveform having a frequency of 1 Hz (Fig 3(c)) was considered in accordance with the literature ^38^. The axial component (1.3 N) of ground reaction force exerted during physiological loading conditions was obtained from the literature ^44^. The proximal end of the tibia was fixed, and the load was applied at the distal end, as shown in Fig 3(a). In accordance with the literature, the periosteal and endocortical bone surfaces were assumed to be completely impermeable and permeable, respectively ^39 38 40^ implemented through zero-flow and zero-pressure boundary conditions (Fig. 3(b)).

**Fig. 3.**
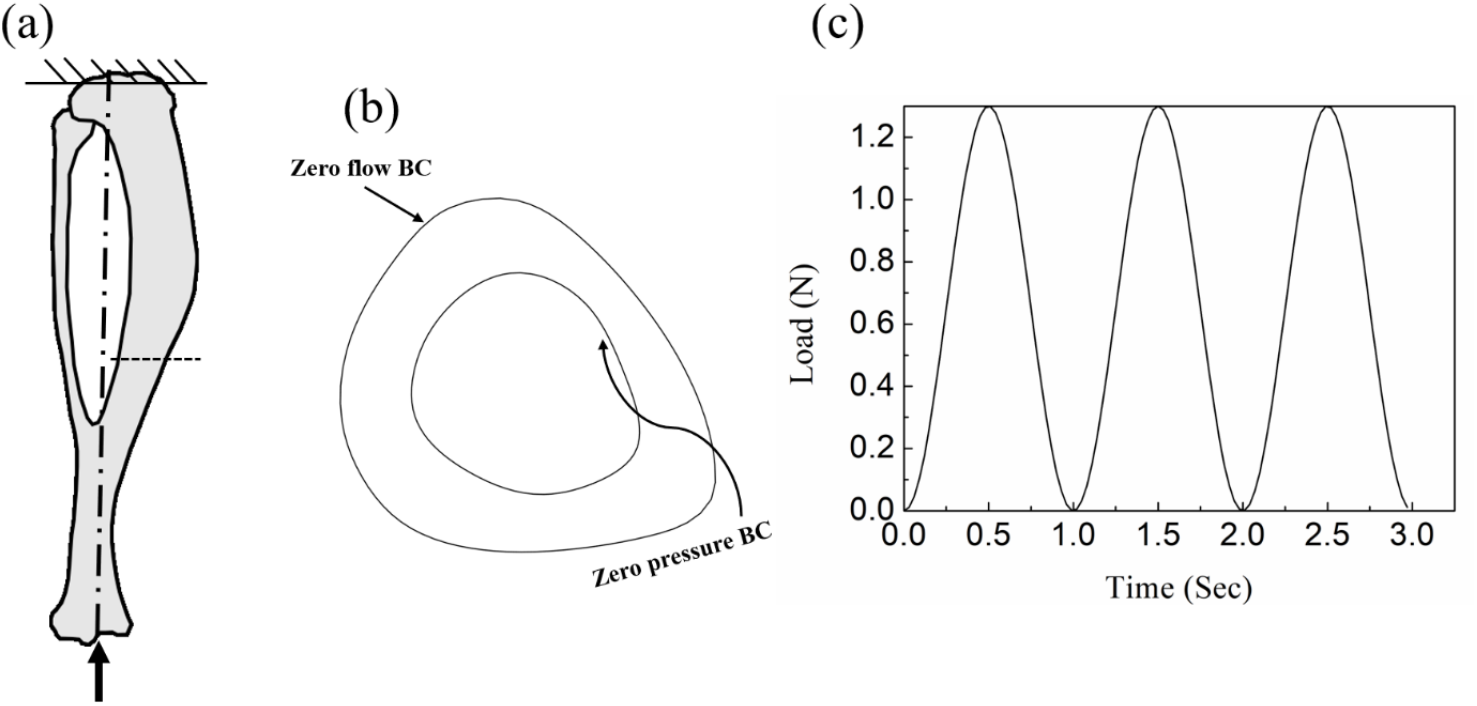
Loading and boundary conditions (a) The proximal end of the tibia was fixed and the distal end was axially loaded with 1.3 N, (b) periosteal and endocortical surface of the bone was prescribed zero-fluid-flow and zero-pressure (*p* = 0) conditions, respectively. (c) To mimic the physiological loading, a Haversine loading waveform of 1 Hz frequency was assumed.

### 2.6 Finite Element Analysis

The finite element problem was conducted using the commercial software ABAQUS (Dassault System). To simulate fully or partially saturated fluid flow through porous media, the soil element (C3D8RP, continuous three-dimensional eight nodded reduced integration pore pressure hexahedral element) type was assigned to the brick element of tibial bone, which is available in ABAQUS ^42^. These elements had 4 degrees of freedom (3 for the displacement and 1 for pore pressure) per node. The bone material was modeled as a linear isotropic poroelastic material whose parameters were taken directly from Kumar et al. ^34^ and have been shown in Table 1. A coupled pore fluid diffusion stress analysis was performed in ABAQUS to capture the fluid flow. The fluid velocity was computed at the integration point during the applied loading. The strain induced by physiological loading (idealized as an axial loading) and corresponding fluid flow velocity at three sections (studied by Ausk et. al. ^20^) viz., distal, mid-section and proximal sections along the tibial length are presented in Fig 4. The strain patterns obtained were also validated against existing literature ^48^.

**Fig. 4.**
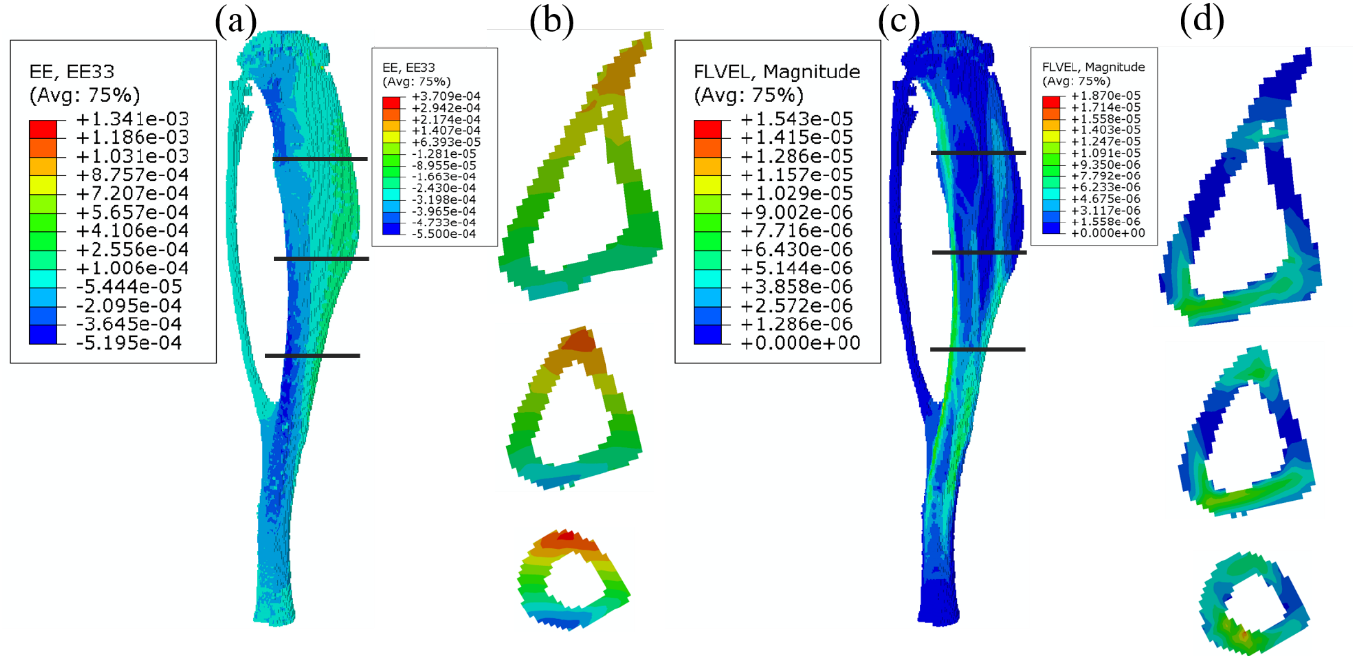
Induced strain and fluid velocity for axially loaded tibia: (a) longitudinal strain contour of a mouse tibia (b) longitudinal strain (EE33) pattern at the three sections, i.e., at distal, mid-section, and proximal (studied by Ausk et al. ^20^) (c) corresponding fluid velocity (mm/s) at the full tibia (d) fluid velocity (mm/s) at the three sections viz. distal, mid, and proximal.

### 2.7 Prismatic Bar Analysis

Creating a full 3D model of as mouse tibia is resources-intensive in terms of software and computational time. The poroelastic analysis of the whole voxel-based tibia under axial loading took 6 hours. In contrast, analyzing a simplified prismatic bar (length 3 mm) under the same conditions reduced the computation time by approximately 80%. Therefore, the tibia was idealized as a prismatic bar with a cross-section similar to that of a C57BL6J, 16-week young adult mouse studied in section 2.6. Three prismatic bars (each with a length of 3 mm) along the tibial length are shown in Fig 5.

**Fig. 5.**
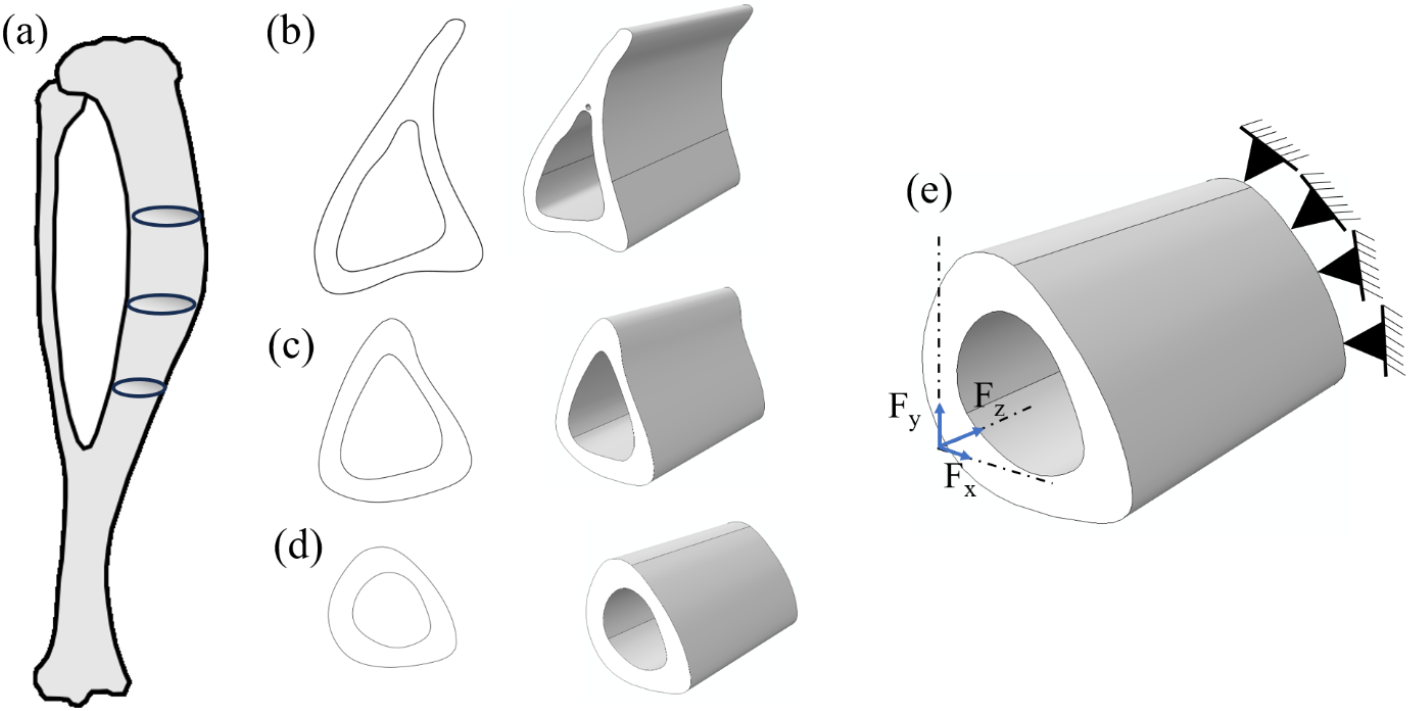
The bone (a) is idealized as a prismatic bar (b) of length 3 mm and having a cross-section similar to that of the “proximal section,” (c) “mid-section” and (d) “distal section” (e) loading condition of a prismatic bar; one end was fixed, and another end was loaded such that it induced strain as in case of a full tibia.

The prismatic bars were meshed with 10650 hexahedral elements after performing a convergence analysis. These bars were modeled as linear poroelastic material, with the material properties kept the same as those of the whole tibia, as outlined in Table 1. One end of each bar was fixed, while the other end was subjected to loading in all three directions (x,y, and z), as shown in (Fig 5(e)), to generate strain similar to the whole tibia. The resulting strain patterns and fluid velocities are shown in Fig 6(b) and Fig 6(c). A comparison of the induced strain and corresponding fluid velocity between the voxel-based 3D tibia and model and the prismatic bar analysis is provided in Table 2.

**Table 2:** Strains and fluid velocities at three distinct sections along the length of a mouse tibia (All strain values are presented in microstrain while fluid velocities are measured in mm/s)

|  | Peak tension surface |  |  | Peak compression surface |  |  |
| --- | --- | --- | --- | --- | --- | --- |
|  | Distal | Mid | Proximal | Distal | Mid | Proximal |
| **Voxel-based 3D tibia**<br>(strain) | 360 | 275 | 189 | -520 | -425 | -277 |
| **Prismatic Bar**<br>(strain) | 354 | 221 | 172 | -541 | -444 | -287 |
|  | Maximum |  |  | Minimum |  |  |
|  | Distal | Mid | Proximal | Distal | Mid | Proximal |
| **Voxel-based 3D tibia**<br>(fluid velocity) | 1.86e-5 | 1.24e-5 | 9.91e-6 | 0 | 0 | 0 |
| **Prismatic Bar**<br>(fluid velocity) | 1.77e-5 | 1.20e-5 | 9.31e-6 | 0 | 0 | 0 |

**Fig. 6.**
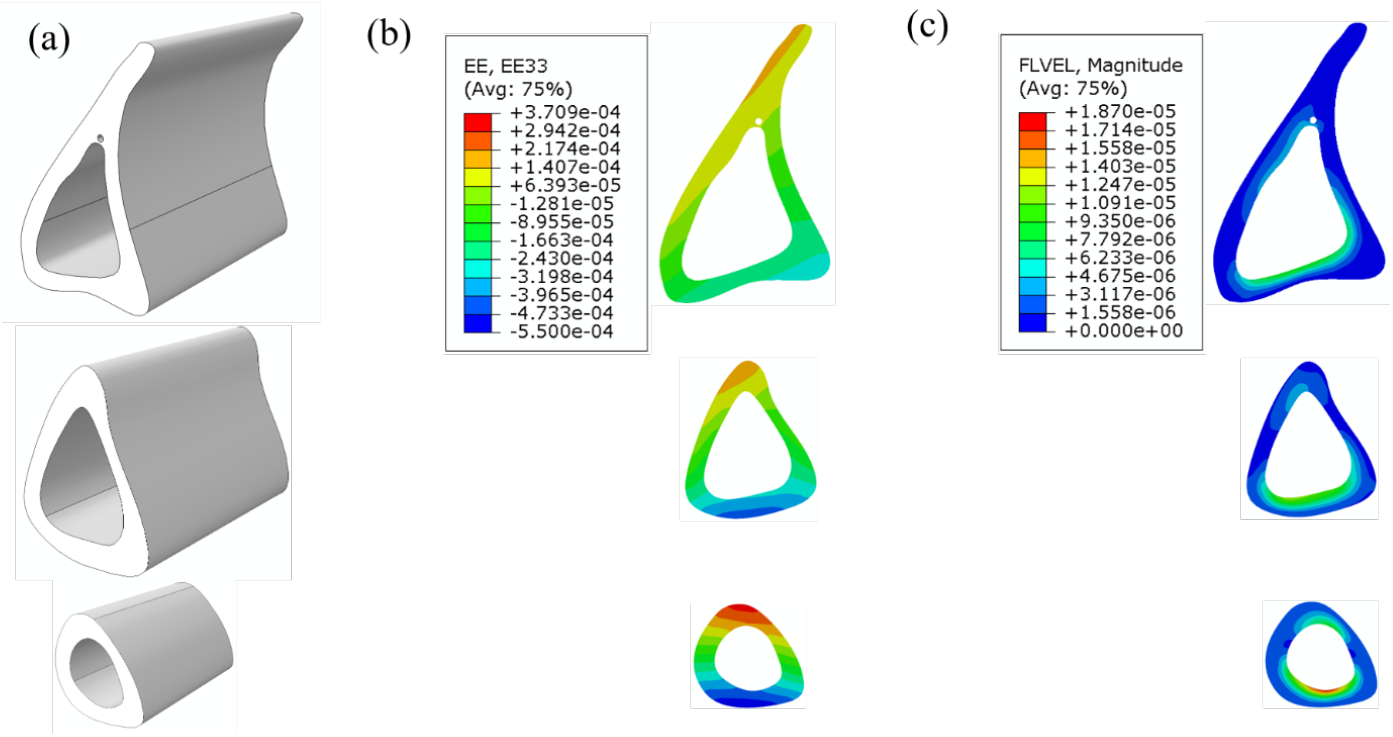
The bone is idealized as a (a) prismatic bar of length 3 mm and having a cross-section similar to that of the “proximal section,” “mid-section,” and “distal section” of an original tibia studied in section 2.6 (b) induced strain at three sections (c) corresponding fluid velocity(mm/s)

### 2.8 Botox-induced disuse Activity

Botox-induced calf muscle paralysis incapacitates the muscles to applying force, which causes a loss of cycling loading on the mouse tibia. Cycling loading is one fundamental stimulus^46^ that causes fluid to flow inside the lacune canalicular channel. We assume that the application of Botox leads to zero fluid velocity inside the porosity of LCN.

### 2.9 Computation of fluid flow-based dissipation energy density

Load-induced viscous fluid flow inside LCN porosities exerts shear stress on the osteocyte cell membrane, and energy is dissipated. The dissipated energy influences the mechano-sensing cell osteocyte, and further biochemical signalling pathways get activated. According to Kumar et al. ^34^, the dissipation energy density (*DED*), which is a function of fluid velocity, can be calculated as:

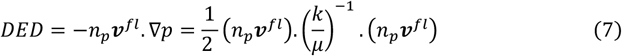

Where *n*_*p*_is the LCN porosity, 𝒱^*fl*^ is the fluid velocity.

### 2.10 Zone of influence

Osteocytes form an interconnected network known as the lacuna-canalicular network (LCN), through which biochemical signals are transmitted. These signals act as a messenger for communication with the other cells, such as osteoblasts, bone lining cells, and osteoclasts. Such communication has been implemented in literature by different mechanisms, such as diffusion ^49^ and zone of influence ^34 50 32^. According to the zone of influence, we capture the contribution of each osteocyte that falls under the given zone. Osteocytes present far from the node of interest (where a total stimulus is being calculated), depicted by a red color in Fig.7(a), contribute less than the osteocytes present near the node of interest. This non-local behavior of osteocytes was implemented by providing a weightage function. We used here an exponential weightage function 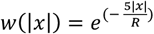 which is per the literature ^5034^ where |*x*| represents the distance between the node of interest and the position of osteocytes, while *R* represents the radius of the spherical zone. For considering the contribution of all the osteocytes, we took R (155 µ*m*), i.e., the radius of the spherical zone equal to the mean of the cortical thickness of the section viz. “distal,” “mid,” and “proximal”. This is in accordance with the earlier work in which optimization was done to get the optimal radius^39^. Considering the ZOI, the dissipation energy density was calculated using Eq. (8) at every i^th^ node of interest. The total dissipation energy density *ϕ* = *DED* ∗ *N* ∗ *d*. Where N is the no. of cycles and d is no. of days.

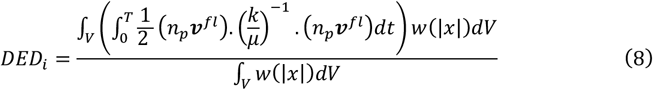

### 2.11 Mineral Resorption Rate (MRR)

Exogenous loading-induced fluid flow inside the LCS has been widely accepted in the literature to correlate with bone adaptation. Recently, Sanjay et al.^50^ have presented a robust formulation that shows that MAR (mineral apposition rate), i.e., rate of change of new bone thickness, is directly proportional to the square root of total dissipation energy density minus its threshold or reference value. That is,

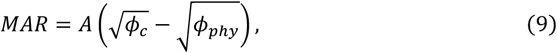

where *A* is the remodelling rate, *ϕ*_*c*_ is the current dissipation energy density, while *ϕ*_*phy*_ represents the dissipation energy density to maintain bone homeostasis (under physiological loading conditions). In the disuse condition, the fluid velocity is zero, which leads 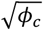 to a zero value. Hence, the expression of MAR is simplified to

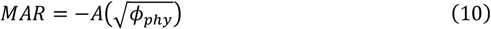

Here, the negative sign shows bone loss, and hence, mineral resorption rate (MRR) can be written as

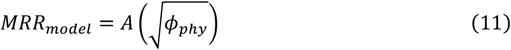

Thus, *MRR*_model_ is directly proportional to the square root of the total dissipation energy density. The parameter *A* is a constant to be determined. The MRR ( *µm*^3^/*µm*^2^/*day*) is defined as the total loss of bone volume per unit of bone surface area per unit time. For the sake of computational convenience, the endocortical bone surface was divided into 360 segments, and then MRR was calculated at every i^th^ point (i.e., a common point of two consecutive segments) as

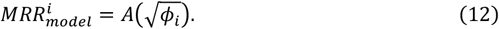

As an example, these points are shown in Fig. 7(b). The constant A was calculated by comparing the experimental bone loss data available in the literature. Experimental endocortical bone loss 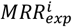 for each of the 360 points was computed from Fig. 7(b) using an in-house MATLAB code. The optimization problem was formulated as a minimization of squared error:

**Fig. 7.**
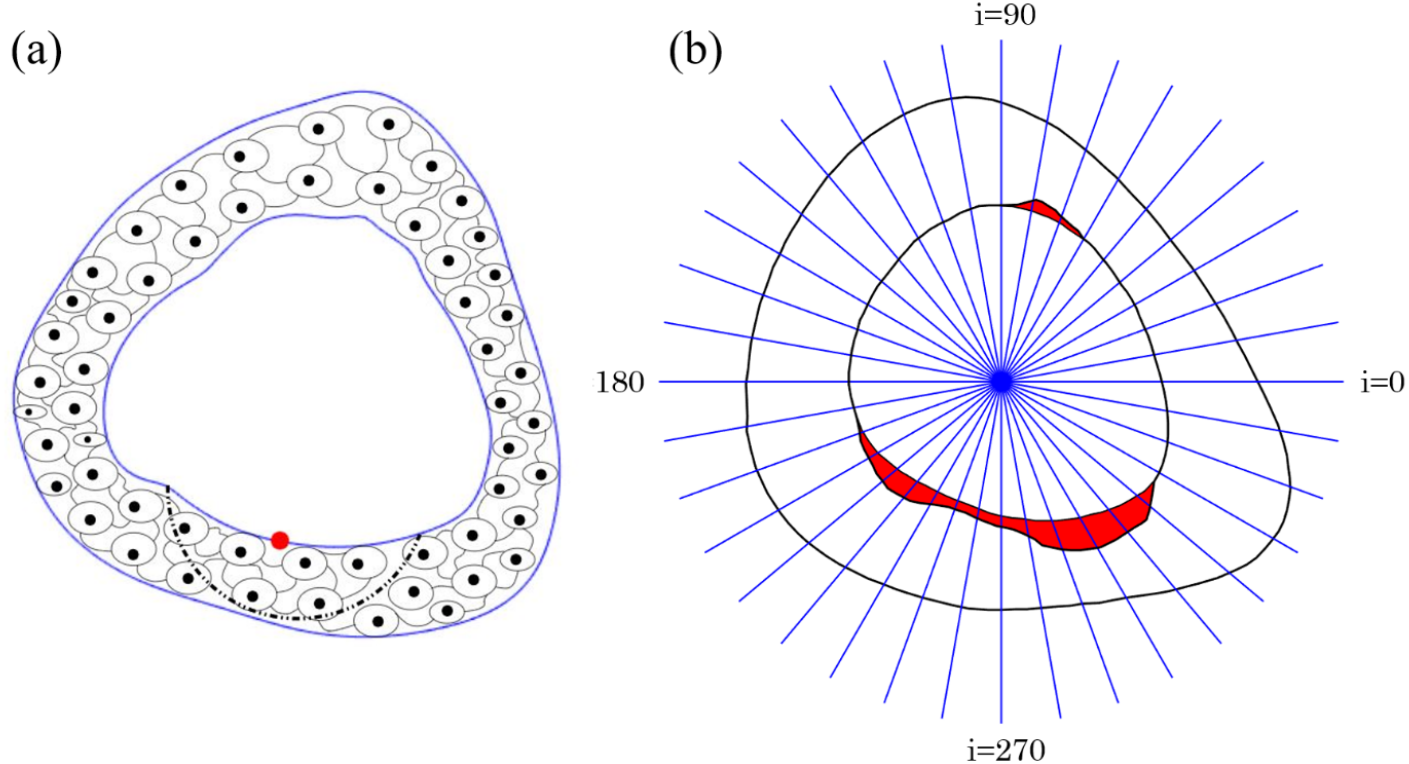
(a) The non-local behavior of the osteocyte is captured through a spherical zone of influence. The red node shows the point of interest. (b) In-vivo cortical bone loss at the distal section of a tibia ^20^; endocortical bone surface was divided into 360 points (for clear visibility, the points have been drawn at every 10 degrees).

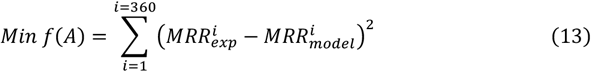

The Levenberg Marquardt algorithm ^51^ available in MATLAB was used to fit the model in Eq. (12) to the experimental data, i.e. 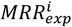 and find the optimal value of the constant *A*. The calculated optimal parameter 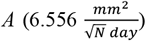 was then used to predict bone loss at the two remaining sections, viz., the mid and proximal sections. A statistical analysis (as detailed in section 2.12) was performed to compare the model prediction to the experimental values.

### 2.12 Statistical Analysis

The endo-cortical bone loss area per unit of time was computed by integrating the MRR over the circumference to compare the model prediction to the experimental values. The bone resorption rate per unit bone surface (BRR/BS) was obtained by dividing the loss area per unit of time by the total perimeter of the surface under consideration (Here, the endocortical surface is under consideration). A one-sample, two-tailed Student’s t-test was used to compare the BRR/BS values. A circular goodness-of-fit test, viz, Watson’s U^2^ test ^52^, was used to compare the endocortical distribution of bone loss predicted by the model to the in-vivo results.

## 3. Results

### 3.1 Distal Section

The developed model was first used to predict cortical bone loss at the distal section. The BRR/BS values have been compared in Fig. 8(d) through a histogram. The obtained site-specific bone loss distribution by the model was shown in Fig. 8(c), whereas the experimental bone loss was shown in Fig. 8(b). The statistical analysis showed that the model BRR/BS, 1.64 *µm*^3^/*µm*^2^/*day*, was not significantly different from the in-vivo BRR/BS, 1.15±0.67 *µm*^3^/*µm*^2^/*day* (p = 0.47, Student’s t-test). The predicted bone loss distribution was also found to be not significantly different from the in-vivo distribution (p = 0.33, Watson’s U^2^ test).

**Fig. 8.**
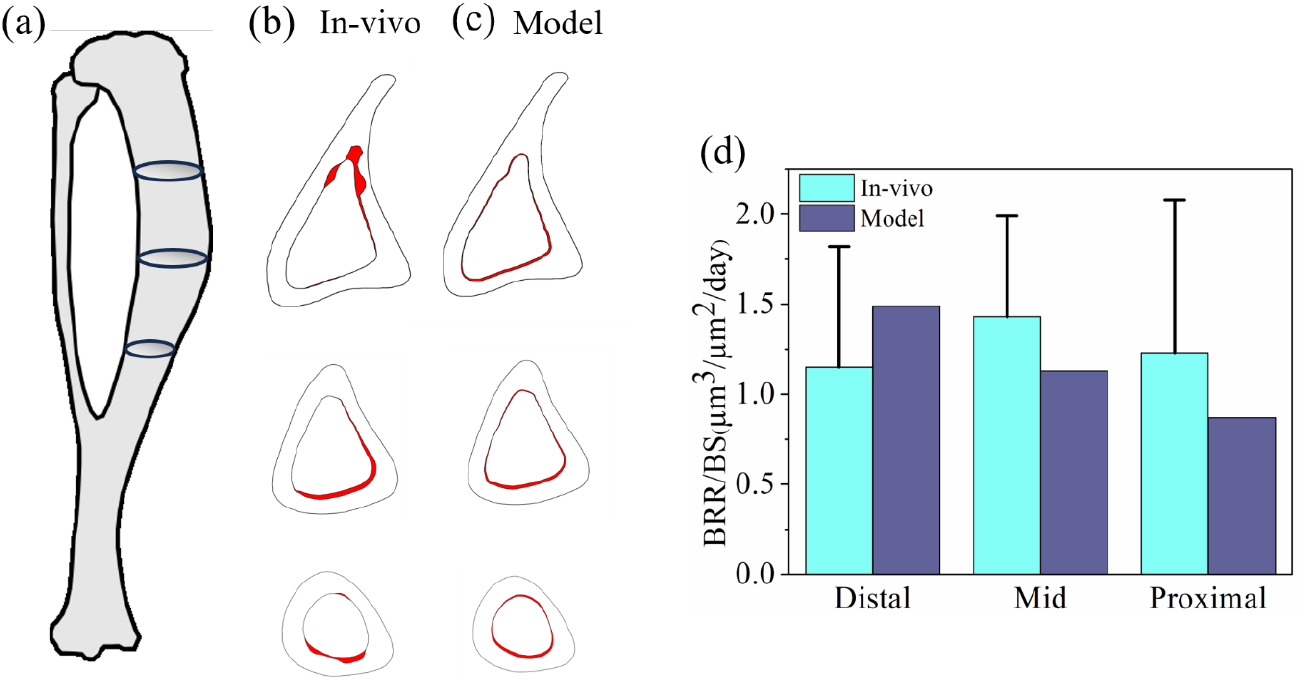
Comparison of the model and the experimental cortical bone loss distribution at the three sections of a mouse tibia: (a) shows the three cross-sections viz. “distal,” “mid,” and “proximal” along the length of a mouse tibia (b) in-vivo, site-specific bone loss ^20^ (shown in red) at the corresponding sections. (c) the bone loss predicted by the model, and (d) a histogram comparing model BRR/BS to the in-vivo value (error bar represents standard error).

### 3.2 Mid-section

Mid-section cortical bone loss was also predicted using the same resorption rate *A*. The predicted and the in-vivo BRR/BS have been compared in Fig. 8(d). The predicted BRR/BS, (1.13 *µm*^3^/*µm*^2^/*day*) was found to be not significantly different from the experimental value of 1.43 ± 0.5675 *µm*^3^/*µm*^2^/*day* (p = 0.61, Student’s t-test). Endocortical bone loss distribution predicted by the model and the in-vivo experiment have also been shown in Fig 8(c) and 8(b), respectively. It was found that the distribution predicted by the model is not significantly different from the in-vivo distribution (*p* = 0.99 for Watson’s U^2^ test).

### 3.3 Proximal section

The bone loss at the proximal section of the same tibial bone was also predicted using the same resorption coefficient *A*. The predicted BRR/BS (0.87 *µm*^3^/*µm*^2^/day) was found to be not significantly different from the corresponding experimental value of 1.23 ± 0.85 (*p* = 0.68, Student’s t-test) as compared in Fig 8(d). The site-specific cortical bone loss distribution predicted by the model (Fig 8(c)) was also found to be significantly different from that of the experimental case (Fig. 8(b)) (*p* = 0.0003 for Watson’s U^2^ test). Hence, the model was not able to predict the experimental bone loss distribution for this case.

### 3.4 Proximal section incorporating muscle attachments

The results so far were obtained based on the axial loading on the tibia as a simplification of the real scenario where different muscles act at different points of time and for different cage activities. This simplification was able to predict the bone resorption for both distal and mid-diaphyseal sections, as detailed in sections 3.1 and 3.2. Since the model was not able to correctly predict the bone loss at the proximal sections, the effect of muscle activity needs to be incorporated. It is evident from the literature that the four significant muscles viz. flexor digitorum longus (FDL), tibialis posterior (TP), extensor hallucis longus (EHL), and tibialis anterior (TA) are attached ^53^ close to the considered proximal section, as schematically shown in Fig. 9(a). As no such muscle attachment is seen close to the remaining sections, viz. the distal and mid-section, analysis of the proximal section may need to be done differently and more accurately. This is also in accordance with the Saint Venant Principle ^54^. Incorporation of muscle activity is complex, however, as there are different cage activities and muscle forces for all these activities are unknown in the existing literature. In view of this, a simple analysis has been carried out just for an example.

**Fig. 9.**
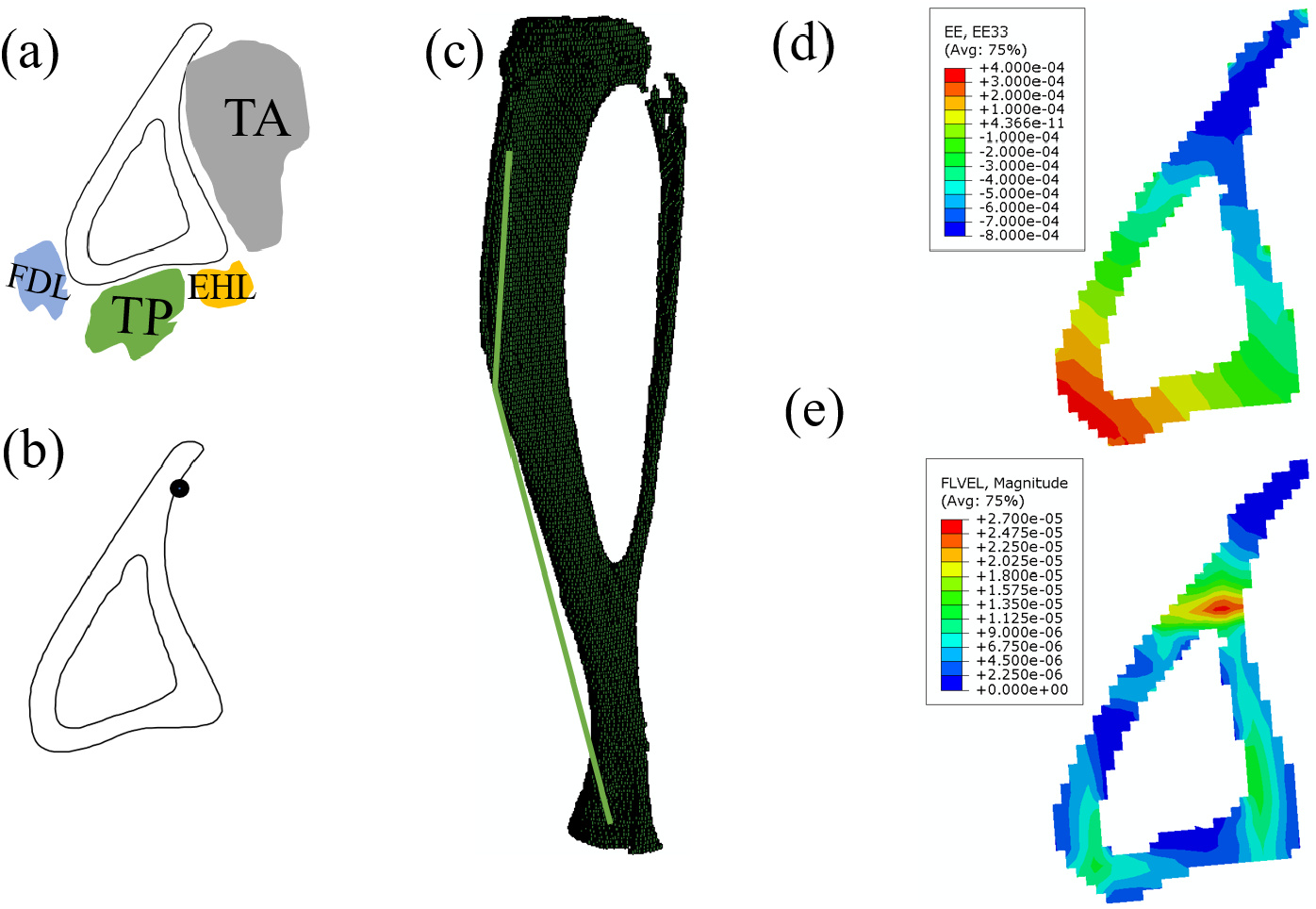
(a) Proximal section with relevant muscles, (b) TA muscle attachment (black dot shows attachment point), (c) shows the TA muscle attachment (through a line diagram) in the whole tibia (d) obtained strain at the considered proximal section (e) corresponding fluid velocity (mm/s).

These muscles attached close to the proximal section of tibia help in dorsi flexion (TA, EHL) and plantar flexion (TP, FDL) of the ankle during walking and, thus apply load on the bone. The maximum load is carried by the dorsi flexion muscle ^55^ during a gait cycle. The dorsi flexion muscle includes tibialis anterior (TA) and extensor hallucis longus (EHL). Between these two muscles, TA is prominent due to its higher physiological cross-sectional area (PCSA); hence, only TA was considered during this study. The attachment point of the TA muscle at the proximal section (Fig. 9(b)) and the corresponding muscle force (2.4 N) were obtained from the literature (Charles et al., 2016). The TA muscle force was applied, keeping the material property and boundary condition the same as earlier. The corresponding strain and fluid flow patterns obtained by the finite element analysis are shown in Figures 9(d) and 9(e), respectively. The experimental and model bone-loss distributions have been shown in Figs. 10(c) and 10(d), respectively. A prismatic bar of the same proximal section was also created and loaded to induce the strain as in the case of the whole tibia, as shown in Fig. 10(a); corresponding fluid velocity was also depicted in Fig. 10(b). The BRR/BS values have also been compared in Fig. 10(e). The predicted BRR/BS (2.07 *µm*^3^/*µm*^2^/day) was found to be not significantly different from the experimental BRR/BS of 1.23 ± 0.85 (*p* = 0.34, Student’s t-test).

**Fig. 10.**
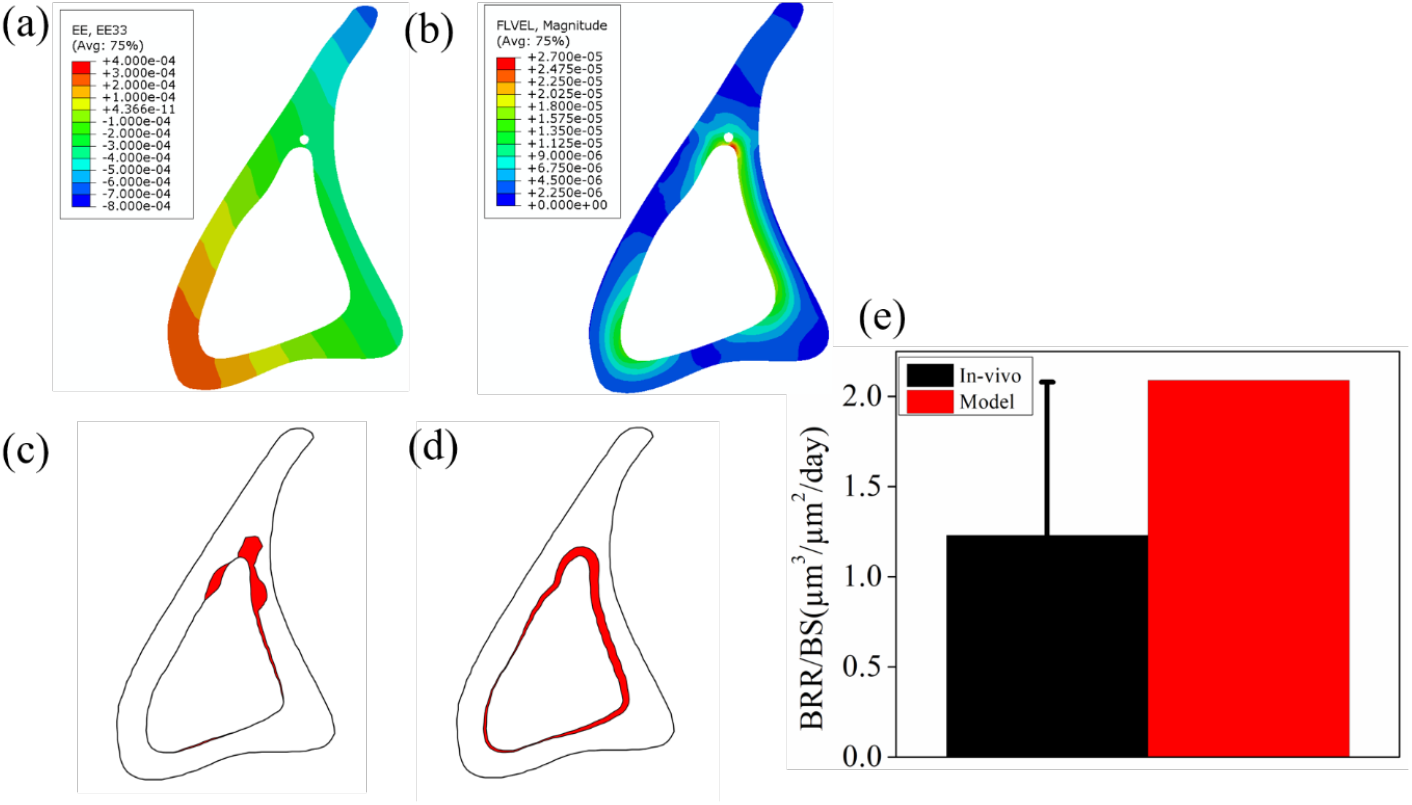
Proximal section with attached muscle idealized as a prismatic bar: (a) Induced strain at the considered proximal section, (b) fluid velocity at the corresponding section (c) in-vivo site-specific cortical bone loss ^20^ (d) site-specific bone loss predicted by the model (e) comparison of model BRR/BS to the in-vivo values.

The bone loss distribution was also found to be not significantly different (*p*= 0.09, Watson U^2^ test). Note that this is only an example showing the importance of incorporating muscle attachments in the current study. However, due to the unavailability of muscle force data for all the cage activity conditions, the detailed analysis incorporating the muscle attachments has been taken as a future work.

### 3.5 Sensitivity Analysis

While developing a mathematical model, it is crucial to analyze how different parameters affect the final output. A sensitivity analysis is necessary to demonstrate how parameter variations influence the result. In this study, we employed a local sensitivity analysis, also referred to as a one-at-a-time sensitivity measure ^57^. This approach involves varying the model parameter by a set amount and observing its impact on the final output. Here, we varied the model parameter remodeling rate (*A*) by a given amount and noted the value of BRR at the sections under consideration, as depicted in Fig. 11. The results indicated that BRR values change linearly with the remodeling rate. Consequently, we identified the remodeling rate (*A*) as a rate of osteocyte apoptosis in the absence of fluid flow, which leads to osteoclast recruitment thus bone loss. This finding aligns with existing literature which identifies a similar parameters associated osteogenesis in response o increased mechanical loading ^50 58^.

**Fig. 11.**
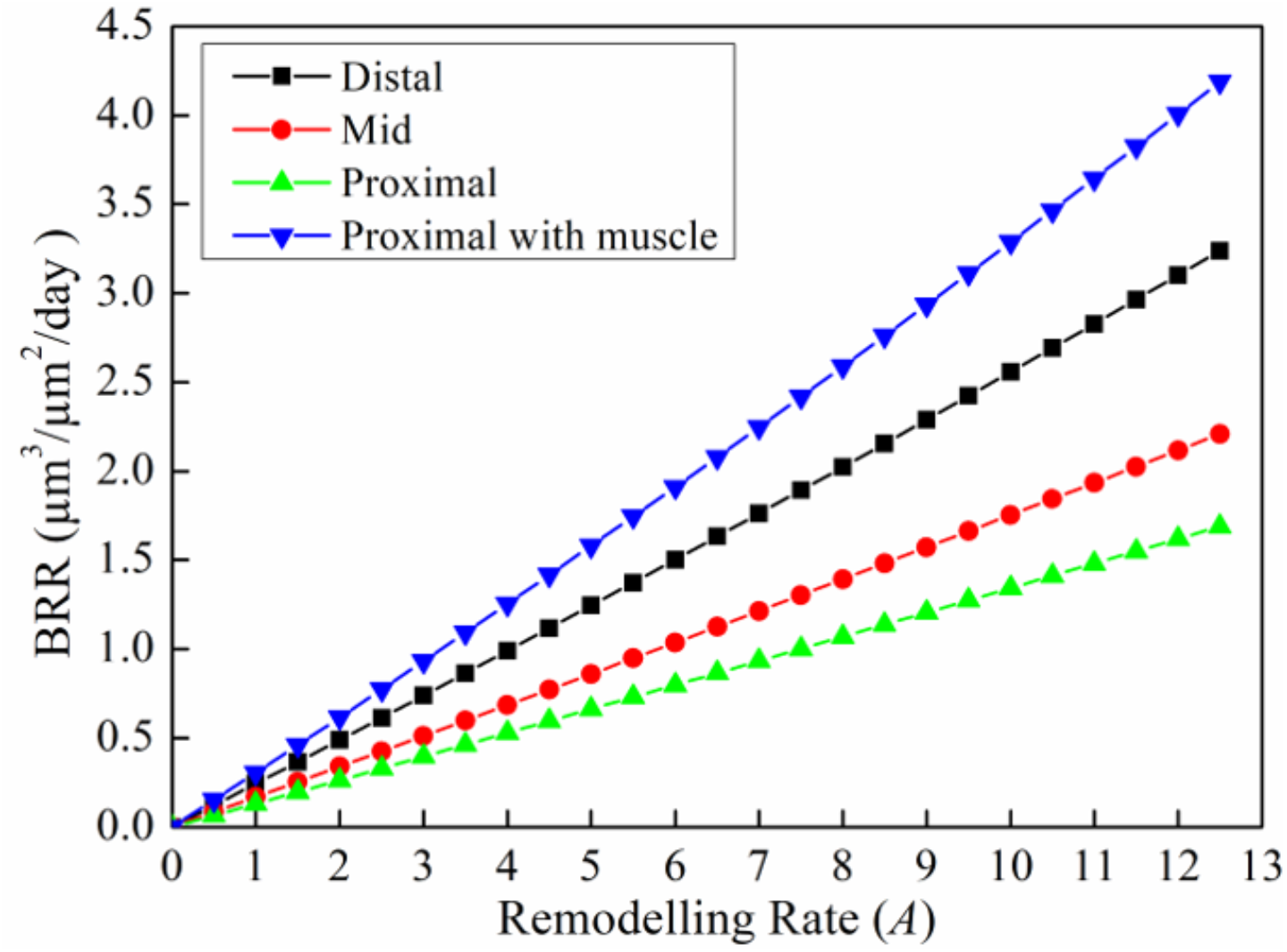
The plot depicts the BRR as a function of remodeling rate (*A*) in the four different sections, viz. Distal, Mid, Proximal, and Proximal with muscle.

## 4. Discussions

The present work studied cortical bone tissue loss due to the disuse of the bone. It developed a fluid flow-based mathematical model that utilized the dissipated energy density as a stimulus. A finite element method was used to simulate fluid flow inside the porosity of LCN, considering bone as a poroelastic material. The model was fitted to the existing bone loss data in the literature. The developed disuse model showed the linear relation between MRR and the 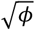 (i.e., the square root of the dissipation energy density), which is in accordance with the earlier work by Sugiyama et al. ^59^. Note that the dissipation energy density is proportional to the square of strain and, hence, MRR being proportional to 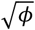 is akin to bone formation rate (BFR) being proportional to the strain ^60^. A work by Singh et al. ^39^ has also shown a similar kind of trend, i.e., the rate of new bone formation is directly proportional to the 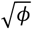. There would be a new bone formation when the stimulus exceeds the threshold value.

It has been widely accepted that osteocytes sense the stimulus for the mechano-transduction. The mechanosensory osteocytes need nutrients for survival, which is achieved via load-induced fluid flow inside the LCS. However, in case of bone disuse, there is no fluid flow, which may incapacitate the flow of nutrients inside the LCS. It may also lead to hypoxia inside the LCN^61^. Such an LCN environment is deleterious for osteocyte viability and may cause apoptosis of osteocytes^62^. Dying osteocytes damage the LCN, which becomes the beacon for osteoclast recruitment ^63 64^. The active osteocytes near the apoptotic osteocytes produce the Receptor activator of nuclear factor kappa beta ligand (RANKL), which binds to the Receptor activator of nuclear factor kappa beta (RANK) present on the surface of osteoclast precursor cells. This initiates osteoclastogenesis, as illustrated in Fig. 12. This theory has been supported by in-vivo studies available in the literature. For example, Emerton et al. showed that mouse ovariectomy caused osteocyte apoptosis and, thus, bone loss via the formation of active osteoclasts ^65^.

**Fig. 12.**
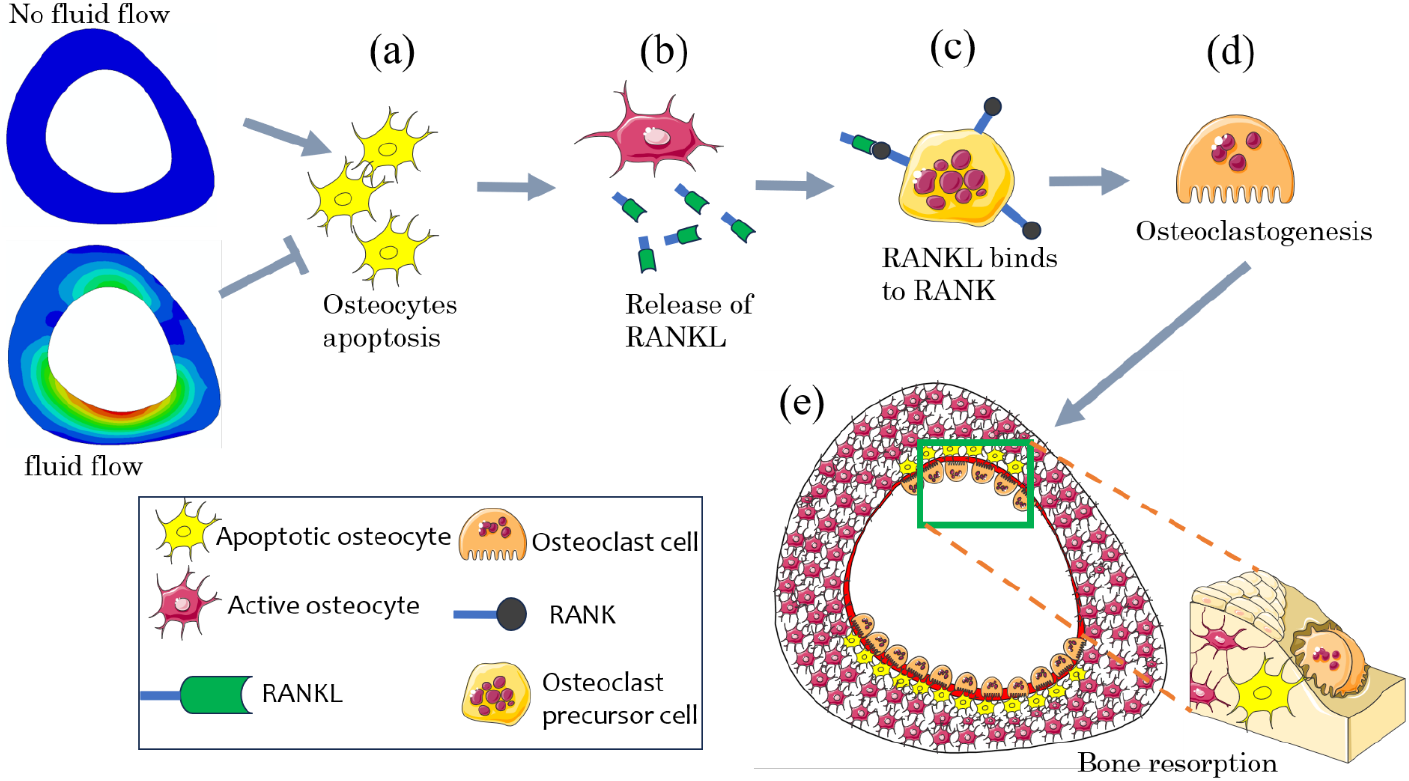
Bone disuse leads to osteoclastogenesis: (a) loss of fluid flow inside the LCS causes apoptosis of osteocytes, (b) active osteocytes near the apoptotic osteocytes release RANKL, (c) RANKL attaches to RANK present on the osteoclast precursor cell, which leads to (d) osteoclastogenesis and (e) bone resorption. (Some parts of this illustration have been created using the Servier medical art, SMART; https://smart.servier.com/)

Another study, in which bone disuse was achieved by a tail suspension model, presented similar trends where bone tissue loss coincided with the site of apoptotic osteocytes ^66^. In a similar study, disuse caused by hindlimb-unloading (via tail suspension) damages the LCN and leads to osteocytic apoptosis resulting into subsequent bone loss ^16^. Although the disuse of bone was achieved via different means, the disuse ultimately caused a loss of stimulus, i.e., a loss of fluid flow inside the LCS. It led to osteocyte-apoptosis and initiated the osteoclastogenesis.

In line with the above studies, the present work has established that the site of maximum bone resorption coincides with that of the maximum stimulus loss, i.e., the maximum loss of dissipation energy density. The developed model predicted cortical bone loss sites at the distal and mid-section of the mouse tibia. However, it could not predict the bone loss distribution at the proximal section. As discussed in section 3.4, this may be due to a different anatomical arrangement (viz. muscle attachment) of the proximal section as compared to that at the distal and mid-diaphyseal sites ^67^.

Exogenous loading-induced bone formation in young-adult mice is seen at both the surfaces viz, periosteal and endocortical surface ^68 69 70 71 72^. Numerous mathematical models have been developed that predict bone formation at the periosteal surface only ^26 49 32^. However, very limited models are available in the literature that predict bone formation at both surfaces ^50^. Initially, tissue strain developed due to increased mechanical loading, which was considered a stimulus for predicting bone formation. However, tissue strain in the whole bone due to physiological loading conditions is 0.2 % ^73^, which is insufficient to elicit the response from the mechanosensory osteocyte. In such a scenario, fluid flows inside the porosity of LCN were explored to amplify the strain required to initiate the intracellular chemical response^7475^. The developed model for increased mechanical loading suggests different stimuli for bone formation at both surfaces. Accordingly, bone disuse should cause stimulus loss and, hence, bone loss at both surfaces. However, in the absence of physiological loading, the bone loss was experimentally found to be at the endocortical surface only ^76 20^. Two different cells are responsible for bone formation and resorption, i.e., osteoblasts and osteoclasts, respectively. It may be possible that, in the case of unloading, these two cells behave differently on the two bone surfaces. Osteoclasts, bone-resorbing cells, may be more pronounced at the endocortical surface in case of disuse than at the periosteal surface due to the easy access to the bone marrow.

Cortical bone loss due to aging was also limited to the endocortical surface only^71^. Skeletal aging lowered the periosteal deposition; however, no resorption was seen at the outer surface.

The earlier study showed that aging reduced the periosteum thickness significantly ^77 78^. The reduced periosteum thickness would, in turn, reduce the pore pressure and thus lower the new bone formation at the periosteal surface, according to the study by Singh et al. ^39^. Modeling age-dependent cortical bone loss has also been taken as a future work.

A study by Gross and Rubin on the turkey’s avian wing bone (viz, radius bone) ^76^ reported the bone loss to be uniform along the length and endocortical perimeter. On the other hand, Ausk et al. ^20^ found that bone loss varies along the length and endocortical perimeter of the mouse tibia. The reason behind this difference may be the difference in structure (curvature, etc.) and loading condition of the two bones (especially torsional loading, which may be prominent in avian bones, as discussed by Novitskaya et al.^79^). Further work is needed to model the turkey’s experimental results. This has also been taken up as a future work.

The developed model predicted the bone loss distribution due to Botox-induced muscle paralysis only. Incorporating the other disuse conditions, such as the tail-suspension model, prolonged bed rest, long space travel, and spinal cord injury, will make the present model robust; however, such experimental data are unavailable in the literature.

Apart from the above, the present model has been developed with several simplifications, as discussed below.

Material properties: The bone is considered a linear isotropic poroelastic material. However, in reality, the bone is highly anisotropic. The cortical bone also has different porosities at different length scales. In the present study, only LCN porosities have been considered. Incorporating all the porosities may enhance the accuracy of the results.

Biological cues: The present model has considered neither bone loss due to natural aging nor the change in the bone’s internal architecture due to aging ^80 81^. Incorporating the aging effect has also been taken into consideration in future work. The present model has not considered the role of different mechanosensory ^21^ such as integrin, tethering fiber, pericellular matrix, etc.

## 5. Conclusions

This work presented a mathematical model for cortical bone loss in disuse conditions. To the author’s best knowledge, there has not been such a model in the literature; hence, the developed model is the first of its kind (however, models are available in the literature for increased mechanical loading). It takes the loss of stimulus due to the disuse of bone as an input and predicts the cortical bone loss at the site of interest. The theory of poroelasticity was used to compute the fluid flow pattern in the LCN, and the corresponding finite element problem was solved using the ABAQUS software. The developed mathematical model predicted the in-vivo bone loss for two murine tibial sites (distal and mid-sections), for which data are available in the literature. However, the model could not predict the bone loss distribution at the proximal section, possibly due to muscle attachment close to this section. A detailed analysis for this section, incorporating muscle attachments on the tibia, may be needed and has been taken as a future work. The current model can be improved by incorporating/predicting other disuse conditions such as extended bed rest, long space travel, and spinal cord injury. However, relevant experimental data will be required for these, which are unavailable in the literature.

## List of Symbols

*σ*_*ij*_: Stress tensor
*ε*_*ij*_: Strain tensor
μ: Viscosity of fluid (Mpa-s)
*p*: Pore pressure (Mpa)
*ν*: Poisson’s ratio
𝒱^*fl*^: Fluid velocity 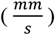
*DED*: Dissipation energy density 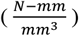
*ϕ*: Total dissipation energy density 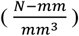
*n*_*p*_: Porosity
BRR/BS: Bone resorption rate per unit bone surface (*µm*^3^/*µm*^2^/day)
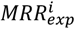: Experimental mineral resorption rate at i^th^ point of interest
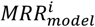: Model mineral resorption rate at the i^th^ point of interest
*q* Fluid flow rate: 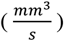
*δ_ij_*: Kronecker delta
*Δ*: Volumetric strain

## Author information

## Contribution

**H.S**. was involved in conceptualization, mathematical modeling, data collection, optimization, model validation, statistical analysis, writing, and manuscript editing. **S.S**. contributed to the software, and **J.P**. supervised and conceptualized the study, contributed to the model review, and edited the manuscript.

## Data availability

The data, such as loading condition, cortical bone loss data, and shape of the bone sections have been collected from the publically available literature as mentioned in the manuscript.

## Ethics declarations

